# Gum Arabic promotes oxidation and ester hydrolysis

**DOI:** 10.1101/199711

**Authors:** Koen P. Vercruysse, Lauren E. Tyler, Jade Readus

## Abstract

We studied the interactions between gum Arabic and select catecholic compounds like caffeic acid, chlorogenic acid and catechol. We observed that GA is capable of promoting the auto-oxidation of the above-mentioned compounds into darkly colored pigments without the addition of redox-sensitive cations. Gum Arabic appeared to be unique among polysaccharide-based materials as many other types of polysaccharides promote the oxidation of the above-mentioned compounds only in the presence of redox cations like Fe^2+^ or Cu^2+^. RP-HPLC and SEC chromatographic techniques were employed to monitor the reactions and to observe the formation of high molecular mass, pigmented materials from gum Arabic and all three catecholic compounds. FT-IR spectroscopic analysis revealed that, despite their darkly colored appearances, the gum Arabic/pigment materials synthesized contain mostly gum Arabic and very little pigment. As chlorogenic acid is an ester of caffeic acid, we studied the capacity of gum Arabic to promote ester hydrolysis using acetylsalicylic acid as the model compound. We observed that gum Arabic did promote the hydrolysis of acetylsalicylic acid into salicylic acid. However, in all our experiments involving the pigment formation between gum Arabic and chlorogenic acid, we did not observe any evidence that chlorogenic acid was hydrolyzed leading to the release of caffeic acid during these reactions. In addition, we observed that heat treatment of gum Arabic did not affect its pro-oxidizing capability, but it did negatively affect its capability to hydrolyze salicylic acid. Thus, these two types of chemical reactivity present in the gum Arabic material may be associated with different components of the gum Arabic material.

## 1. Introduction

Gum Arabic (GA) is described as the dried exudation obtained from the stems of closely related species of Acacia.[1] GA has found applications as stabilizer, thickening agent and emulsifier; particularly in the food industry.[2] The chemical composition of GA is complex, containing mainly polysaccharide (PS) and some protein.[3] The composition of GA changes depending on the exact species of *Acacia* it is derived from, the age of the tree, climatic and soil conditions.[1]

In a previous report, we described the effect of various PS on the oxidation of catecholamines (CAs) into melanin (MN)-like pigments.[4] That report did not include our observations regarding the effects of GA on the oxidation of CAs. It appeared that, compared to the other PS materials tested, GA had only a modest effect on the MN formation from CAs. However, these results were expanded by evaluating the effect of PS and GA on the oxidation of other catecholic compounds. In the course of these experiments, we observed that GA did have a marked effect on pigment formation from compounds like caffeic acid (**(1)** in Fig. 1), chlorogenic acid (**(2)** in Fig.1) or catechol (aka 1,2 dihydroxybenzene; **(3)** in Fig.1). Many of the PS that readily promoted the oxidation of CAs into MN-like pigments, appeared to have only a weak or modest effect on the oxidation of **(1)**, **(2)** or **(3)**. Thus, this report focuses on the apparent unique interactions between GA and select phenolic compounds like **(1)**, **(2)** or **(3)**.

As in other kingdoms of life, plants are capable of producing MN-like pigments and **(1)** or **(3)**, together with other polyphenols, can serve as precursors for such types of MNs; then sometimes referred to as allomelanins.[5] The “browning” in, e.g., cut fruits or vegetables, is due to the formation of MN and is often attributed to the action of enzymes like polyphenol oxidase (PPO), catechol oxidase or tyrosinase.[6] In plants, flavonoids, another class of phenolic compounds, can be converted into pigments known as tannins. In fact, there are similarities in the biosynthesis and biochemistry of MNs and tannins.[7] Such comparisons can be extended to another class of plant-derived pigment: the lignins. Lignins are polymeric substances made from select phenolic precursor molecules.[8] In the case of the biosynthesis of lignins, a recent study emphasized the importance of the hydrolysis of **(2)**, an ester of **(1)**, by the enyzme caffoyl shikimate, aka chlorogenic acid esterase (CSE).[9] Thus, in parallel with our study of the oxidation of **(1)** and **(2)** in the presence of GA, we studied the potential of GA to promote the hydrolysis of esters. For comparison purposes, we studied the hydrolysis of acetylsalicylic acid (**(6)** in Fig. 1) to salicylic acid (**(5)** in Fig.1) in the presence of GA. **(5)** is a plant-based phenolic compound, but preliminary experiments had indicated that both **(5)** and **(6)** appeared not to generate any pigmented materials when incubated with GA or any other PS materials.

**Fig. 1:**
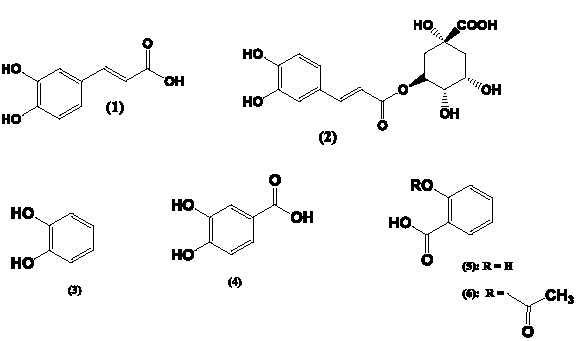
Chemical structures of compounds involved in this study. **(1)** = caffeic acid**; (2)** = chlorogenic acid; **(3)** = catechol; **(4)** = 3,4-dihydroxybenzoic acid; **(5)** = salicylic acid; **(6)** = acetylsalicylic acid.

## 2. Experimental section

### 2.1 Materials

Gum Arabic, caffeic acid, chlorogenic acid, catechol, 3,4- dihydroxy benzoic acid, acetylsalicylic acid, salicylic acid, chondroitin sulfate type A (sodium salt from bovine trachea; 70% with counterbalance chondroitin sulfate type C), chondroitin sulfate type C (sodium salt from shark cartilage; 90% with counterbalance chondroitin sulfate type A), alginic acid sodium salt (Algin®, sodium alginate) - carrageenan (commercial grade, type II), carboxymethylcellulose (low viscosity grade; sodium salt), dextran sulfate, fucoidan (from *Fucus vesiculosus*) and L-ascorbic acid were obtained from Sigma-Aldrich (Milwaukee, WI). Sodium acetate and CuCl_2_.2H_2_O were obtained from Fisher Scientific (Suwanee, GA). All other reagents were of analytical grade.

### 2.2 Stock solutions and reaction mixtures

Stock solutions of CuCl2.2H2O were prepared in water in advance. Stock solutions of GA or PS were prepared in water one day prior to the start of the experiments to ensure proper dissolution. The pH of these GA or PS solutions were measured and typically ranged between 6.4 and 7.5. Stock solutions of compounds **(1)** to **(6)** were prepared just prior to the start of the experiments by dissolving the compounds in methanol and adding water such that the final methanol concentration was 10% v/v.

### 2.3 UV/Vis spectroscopy

Kinetic UV/Vis spectroscopic analyses were performed in 1mL cuvettes using a DU 800 spectrophotometer from Beckman Coulter (Fullerton, CA) against water as the blank. The absorbance at 400nm was measured as a function of the reaction time and all experiments were performed at room temperature (RT).

Microwell UV/Vis spectroscopic measurements were made using the SynergyHT microplate reader from Biotek (Winooski, VT). Aliquots of 200μL of reaction mixtures or purified material solutions were placed in the wells and 200μL water was used as the blank.

### 2.4 RP-HPLC analyses

RP-HPLC analyses were performed on a UFLC chromatography system equipped with dual LC-6AD solvent delivery pumps and SPD-M20A diode array detector from Shimadzu, USA (Columbia, MD). Analyses were performed on a BDS Hypersil C8 column (125X4.6 mm) obtained from Fisher Scientific (Suwanee, GA). Area-under-the-curve (AUC) data were obtained using the LCSolutions software. Analyses of samples containing **(1)** or **(2)** were performed in an isocratic fashion using a mixture of water:methanol:acetic acid (90:10:0.05% v/v) as the solvent and the sample volume was 20μL. Samples were diluted with RP-HPLC solvent such that the concentration of **(1)** or **(2)** in the aliquots from the control experiments was 0.1mM.

Analyses of samples containing **(5)** and **(6)** were performed in an isocratic fashion using a mixture of water:methanol:acetic acid (70:30:0.05% v/v) as the solvent and the sample volume was 20μL. Samples were diluted with RP-HPLC solvent such that the concentration of **(5)** or **(6)** in the aliquots from the control experiments was 0.1mM.

### 2.5 Size exclusion chromatography (SEC)

SEC analyses were performed as described elsewhere.[4]

### 2.6. FT-IR spectroscopy

FT-IR spectroscopic scans were made as described elsewhere.[4]

## 3. Results

### 3.1 MN formation from (1) and (2)

#### 3.1.1 UV_Vis spectroscopy studies

A set of kinetic experiments were performed to evaluate the effect of GA on the oxidation of **(1)** in the absence or presence of Cu^2+^. At a final concentration of 1.0mM **(1)** was mixed with either: (a) water, (b) Cu^2+^ (final concentration arbitrarily fixed at 0.4mM), (c) GA (final concentration arbitrarily fixed at14 mg/mL) and Cu^2+^ (final concentration 0.4mM) or (d) GA (final concentration14 mg/mL). As the spectrophotometer used for these kinetic studies had six cuvette positions, the experiment containing the GA and Cu^2+^ mixture was performed in triplicate. Fig. 2, panel A, shows the kinetic profiles, recording the absorbance at 400nm as a function of the reaction time, thus obtained.

**Fig. 2:**
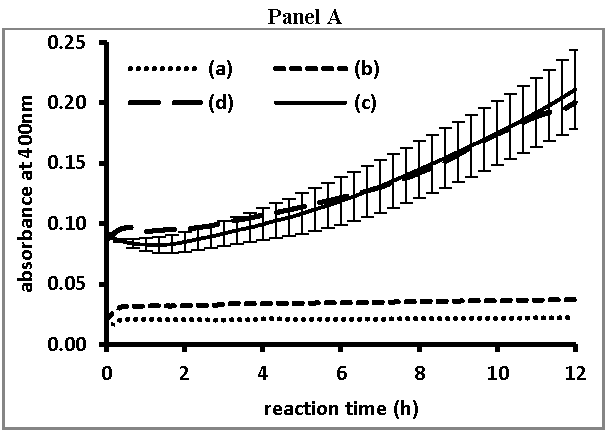
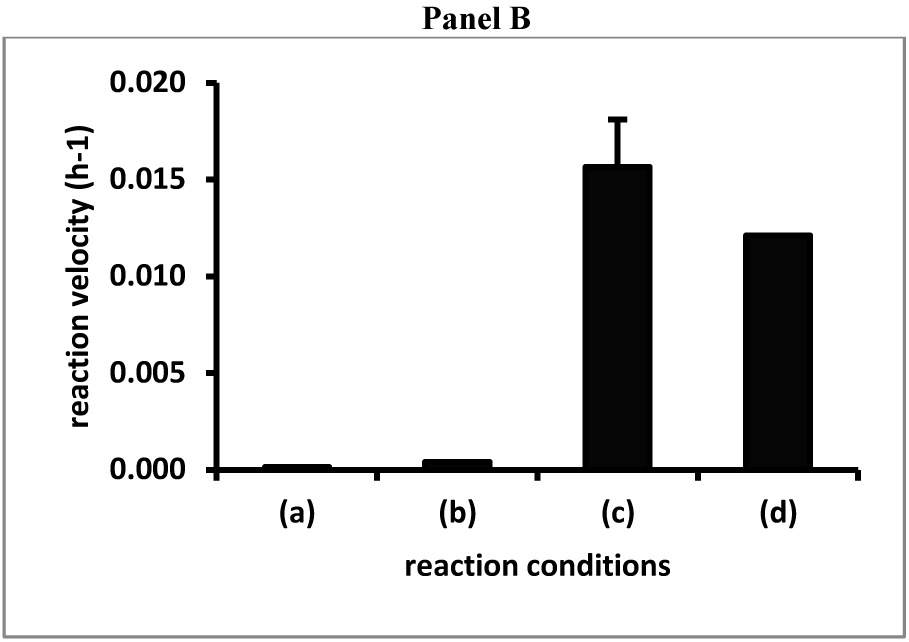
Kinetic profiles of the interaction between GA and **(1)** in the presence or absence of Cu^2+^. **Panel A** presents the absorbance at 400nm as function of the reaction time for mixtures containing 1mM **(1)** and (a) water, (b) Cu^2+^ (0.4mM), (c) GA (14 mg/mL) and Cu^2+^ (0.4mM) or (d) GA (14 mg/mL) The reaction involving the combination of GA and Cu^2+^ was performed in triplicate and the average absorbance ± standard deviation (n=3) are shown. **Panel B** compares the velocities (slopes of the linear regression between 3 and 12 hours) of the kinetic experiments shown in Panel A. The reaction involving the combination of GA and Cu^2+^was performed in triplicate and the average slope ± standard deviation (n=3) is shown.

The kinetic profiles show that the absorbance at 400nm did not increase until about 2 hours into the reaction and then increased linearly as a function of the reaction time. For each reaction profile, a linear regression was performed between 3 and 12 hours of reaction and the slopes thus obtained were taken as a measure of the velocity of the reaction. Fig. 2, panel B, shows a comparison of these slopes for the reactions involving **(1)** in the absence or presence of Cu^2+^ and/or in the presence of GA. For the reaction involving **(1)**, GA and Cu^2+^, the average slope ± standard deviation of the three reactions was determined and shown in Fig. 2, panel B.

Overall, the results presented in Fig.2 show that GA enhanced the oxidation of **(1)**, independent of the presence of Cu^2+^. Many PS, e.g., chondroitin sulfate (CS) A or C, that were capable of promoting the oxidation of CAs[4], were capable of promoting the oxidation of **(1)**, but not without the presence of transition elements like Cu^2+^ or Fe^2+^ (results not shown).

Mixtures containing **(1)** (final concentration 1mM) were mixed with GA (final concentration ranging between 0 and 20mg/mL) and left at RT overnight. All reaction mixtures were set up in triplicate. Aliquots of the mixtures were transferred to wells of a microplate and the absorbance at 400nm was measured against water as the blank. Fig. 3 provides the relationship between the concentration of GA present and the average absorbance at 400nm.

**Fig. 3:**
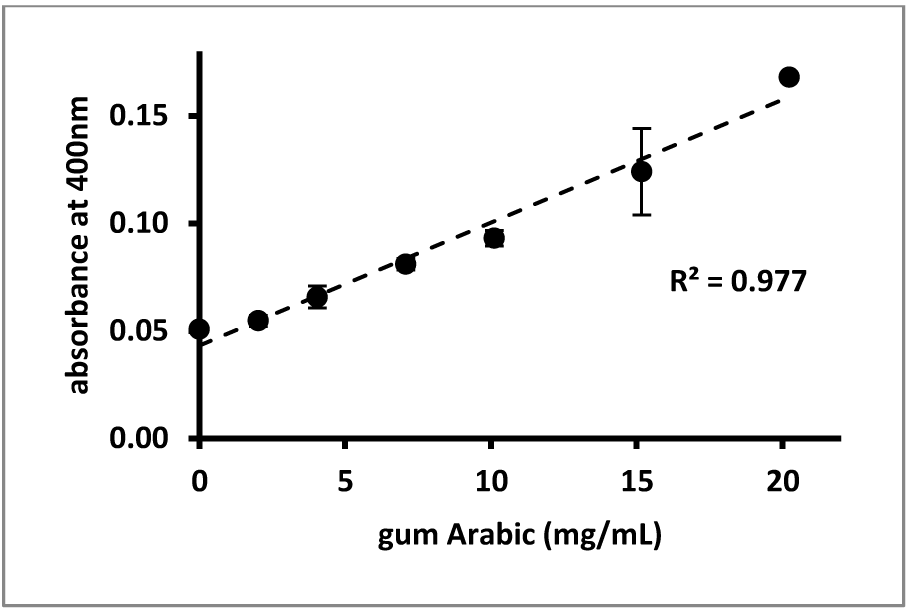
Relationship between absorbance at 400nm and GA concentration for reactions involving **(1)**. Average absorbance at 400nm ± standard deviation (n=3) of mixtures containing 1mM **(1)** and between 0 and 20mg/mL GA. All reaction mixtures were set up in triplicate and left at RT overnight.

The results presented in Fig. 3 indicate that the absorbance at 400nm increased linearly as a function of the GA concentration. Although the reaction proceeds well at RT, increasing the temperature to 37°C or higher increased the velocity of the reaction with changes of color visible within hours upon starting the reaction (results not shown).

A similar pattern of results were obtained for reactions involving GA and **(2)** (results not shown).

### 3.1.2 RP-HPLC studies

As for our studies involving the oxidation of CAs in the presence of PS [4], RP-HPLC allowed for the analysis of **(1)** (retention time about 6.8min) or **(2)** (retention time about 4.5min), but could not provide a resolution between GA (retention time less than 1min) and new products (reaction times of about 1min) that emerged during the reactions between GA and **(1)** or **(2)**. However, RP-HPLC analyses were routinely used to evaluate the disappearance of **(1)** or **(2)** during the various experiments.

RP-HPLC analyses were used to evaluate the effect of heat treatment of GA on its capacity to promote the formation of colored pigment from both **(1)** and **(2)**. In these experiments GA solutions were freshly prepared and kept at RT or kept at 60°C for six hours. Reaction mixtures were set up containing **(1)** or **(2)**, final concentration of 1.0mM, and GA, kept at RT or heat treated at a final concentration 7.0mg/mL. Fig.4 presents a comparison of the relative AUC (integration using the signal at 325nm) of **(1)** or **(2)** after overnight reaction. In this figure, condition (a) refers to **(1)** or **(2)** solutions kept at RT overnight, condition (b) refers to or **(2)** solutions kept at 37°C overnight, condition (c) refers to **(1)** or **(2)** incubated overnight at 37°C using GA that was kept at RT and condition (d) refers to **(1)** and **(2)** incubated overnight at 37°C with heat treated GA. Relative AUC values were calculated using the AUC of the aliquots from solutions containing **(1)** or **(2)** kept at RT overnight as the reference.

**Fig. 4:**
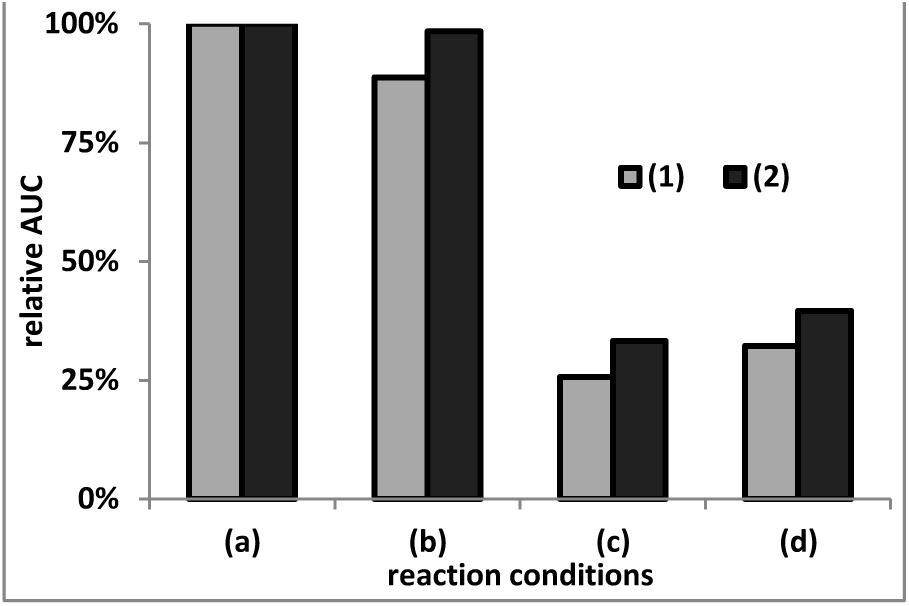
Effect of GA (kept at RT or heat treated) on its reactivity towards (1) or (2). Reaction mixtures containing (1) or (2), final concentration of 1.0mM, and GA, kept at RT or heat treated at a final concentration 7.0mg/mL. Reaction conditions were (a) (1) or (2) solutions kept at RT overnight, (b) (1) or (2) kept at 37°C overnight, (c) (1) or (2) incubated at 37°C using GA that was kept at RT and (d) (1) and (2) incubated overnight at 37°C with heat treated GA. Relative AUC values were obtained using the AUC from reaction condition (a) as the reference.

When incubated at 37°C in the absence of any GA, less than a 10% decline in AUC for both **(1)** and **(2)** was observed compared to the incubation of **(1)** or **(2)** at RT. Incubation with GA resulted in a loss of up to 75% of **(1)** and **(2)**. The pretreatment of GA at 60°C appeared not to have any effect on the outcome of the experiment, although we cannot exclude that the kinetics of the reaction may have been different depending on whether the GA was heat treated or not. Additionally, all the RP-HPLC analyses involving reactions with **(2)** did not show any peak with retention time corresponding to **(1)**. Thus, the reaction between GA and **(2)** did not involve any hydrolysis of **(2)** with release of **(1)** or the reaction between GA and **(2**) did involve hydrolysis, but the thus generated **(1)** immediately reacted further.

#### 3.1.3 SEC studies

Figs. 5 and 6 present some SEC profiles (viewed at 250 or 400nm) for aliquots of the reaction mixtures involving **(1)** or incubated with GA (kept at RT or at 60°C) described in the previous section. Fig. 5, panels A and B, present SEC profiles of GA and **(1)** mixed with GA that was kept at RT after overnight reaction at 37°C. Panel A of Fig. 5 presents the full, 120 minute, SEC profiles, while panel B presents the first 30 minutes of these SEC profiles.

**Fig. 5:**
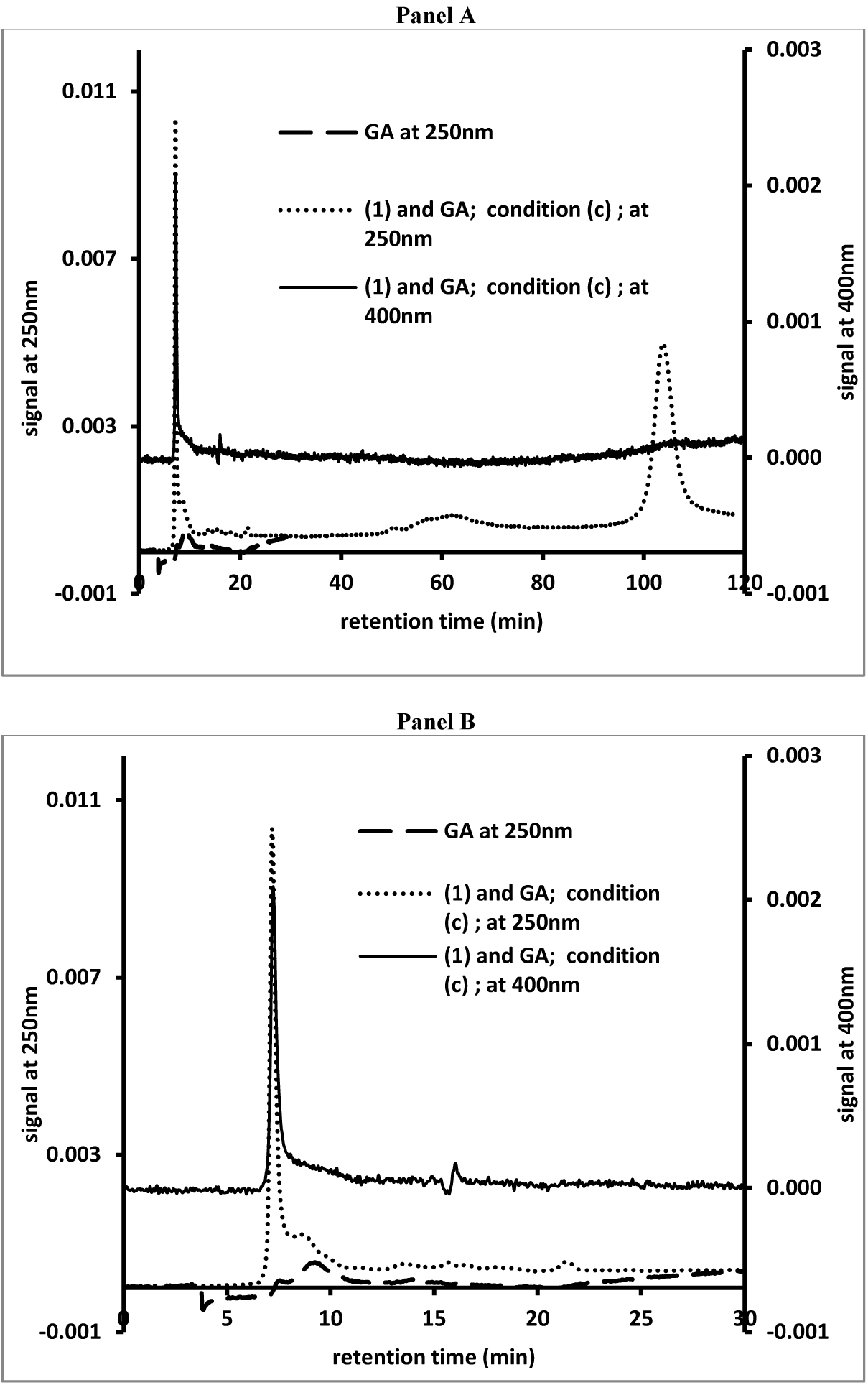
SEC profiles of the reaction mixture containing GA and **(1)**. SEC analysis of GA (viewed at 250nm) and an aliquot from the reaction mixture (viewed at 250 and 400nm) containing **(1)** and GA (condition (c) as described for Fig. 4) after overnight reaction at 37°C. **Panel A** presents the full 120 minute profiles, while **panel B** presents the first 30 minutes of the profiles.

Fig. 6 present similar SEC profiles for GA and **(2)** mixed with GA that was kept at RT. The profiles shown were obtained after two seven of reaction at 37°C.

**Fig. 6:**
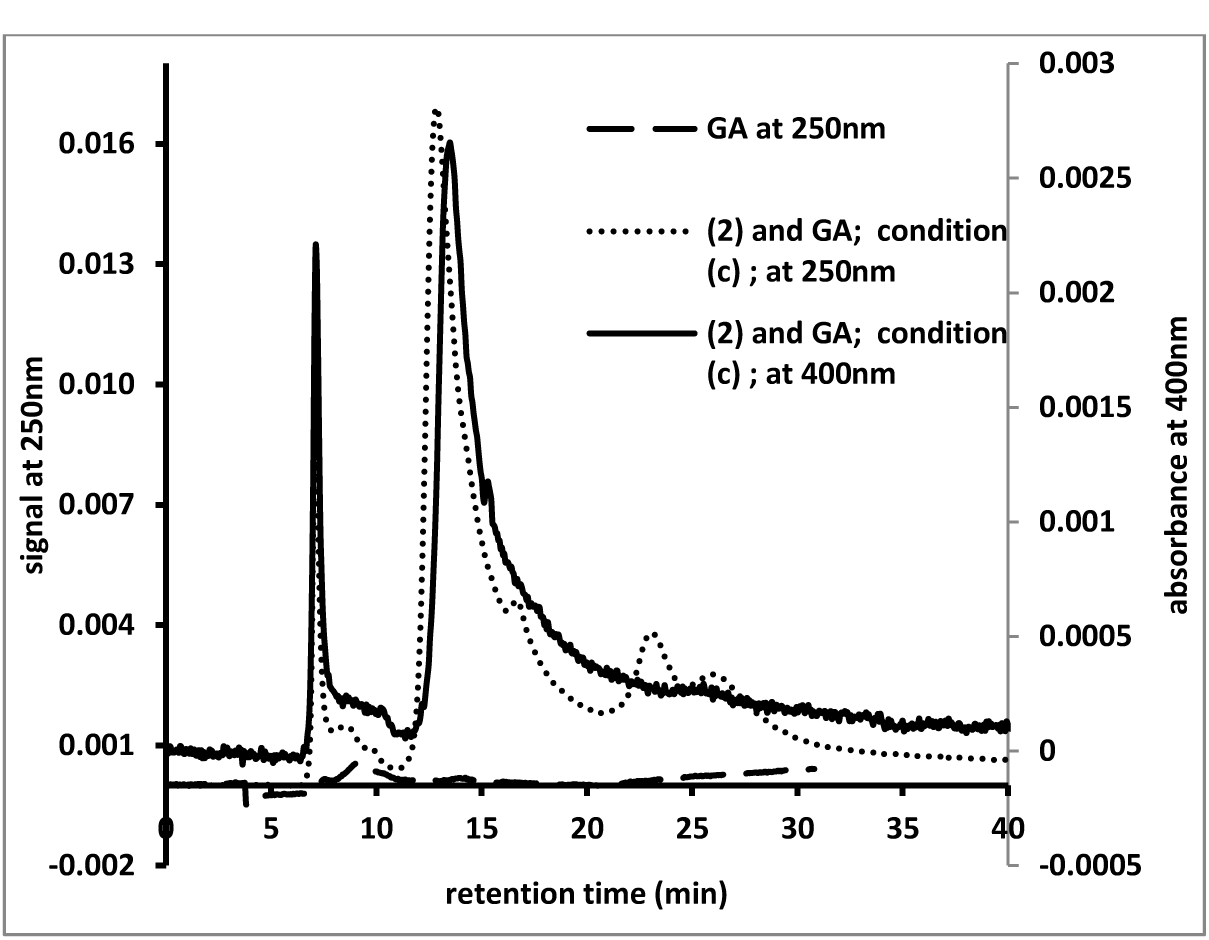
SEC profiles of the reaction mixture containing GA and **(2)**. SEC analysis of GA (viewed at 250nm) and an aliquot from the reaction mixture (viewed at 250 and 400nm) containing **(2)** and GA (condition (c) as described for Fig. 4) after seven days of reaction at 37°C.

For discussion purposes we subdivide the SEC profiles into four different regions. Region I includes peak retention times between 5 and 7 minutes. Such peak retention times are often observed following the injection of PS-stabilized gold or silver nanoparticles^1^ and thought to be close to the upper exclusion limit of the column. Region II includes peak retention times between 7 and 14 minutes. Such peak retention times are typically observed following the injection of GA or other PS solutions and are associated with the size distribution profiles of these materials. Region III includes peak retention times between 14 and 15 minutes. Such peak retention times are observed following the injection of water and thought to represent the lower exclusion limit of the column. Region IV includes peak retention times above 15 minutes. Such retention times are typical for molecules with retentions determined by adsorption phenomena in addition to size-exclusion phenomena.

Fig. 5, panels A and B, show the SEC profiles, of an aliquot from a reaction between GA (stock solution kept at RT) and **(1)** after overnight reaction at 37°C. All reaction mixtures were kept at 37°C for up to seven days and analyzed on a few occasions using SEC. After overnight reaction, the mixture had developed a yellow-orange color. When viewed at 250nm, GA showed a broad peak with little absorbance in region II of the SEC profiles. When viewed at 250nm, **(1)** showed a peak retention time of about 100 minutes, region IV of the SEC profiles. Following reaction with GA, a broad peak in region II of the SEC profiles with little UV_Vis absorbance could be observed. However, a sharp peak emerged in region I of the SEC profiles. The AUC of this sharp peak increased over the course of the multi-day experiment. No other new peaks could be observed. When viewed at 400nm only a sharp peak in region I of the SEC profiles could be observed. The AUC of this peak increased significantly during the course of the multi-day experiment. Thus all the pigmented material generated in this reaction appeared to be associated with materials of very high molecular mass. The reaction between **(1)** and GA, kept at 60°C for multiple hours prior to the start of the reaction, yielded similar results. As observed using RP-HPLC analyses, heat treatment of GA did not affect the reaction between GA and **(1)**.

Fig. 6 shows the SEC profiles of an aliquot from a reaction between GA (stock solution kept at RT) and **(1)**. All reaction mixtures were kept at 37°C for up to seven days and analyzed on a few occasions using SEC. As for the reaction involving **(1)**, the reaction mixture containing **(2)** had developed a yellow-orange color after overnight reaction. SEC analyses of freshly prepared solutions containing **(2)** showed a peak retention time of about 25 minutes in region IV of the SEC profiles. During the course of the reaction, the AUC of the peak corresponding with **(2**) declined and a new peak could be observed close to the peak corresponding with **(2)** in region IV of the SEC profiles. In addition, new peaks emerged with retention times of 13.6 and 17.4 minutes in regions III and IV of the SEC profiles. A broad peak with little absorbance could be observed in region II of the SEC profiles. Following seven days of reaction, the AUC of the peaks observed in region IV had declined, while the AUC of the peaks observed in region III increased. As for the reaction with **(1)**, a sharp peak appeared in region I of the SEC profiles. Although the SEC profiles shown in Fig. 6 indicate a time range of 40 minutes, all the analyses involving **(2)** were run for 120 minutes to evaluate the possibility that hydrolysis of **(2**) with release of **(1)** had occurred. None of the SEC analyses involving the reactions with **(2)** showed any signs of the presence of **(1)**. In Fig. 6, when viewed at 400nm, sharp peaks in region I of the SEC profiles and peaks in region III of the SEC profiles can be observed. The peak in region I was not present after two days of reaction, while the peaks in region III were present after two days of reaction and their AUC increased during the course of the reaction. A similar reaction between **(2**) and GA, kept at 60°C for multiple hours prior to the start of the reaction, yielded similar results. As observed during the RP-HPLC experiments, heat treatment of GA did not affect the reaction between GA and **(2)**.

### 3.2 Large scale reactions

In this set of experiments, the effect of GA on the oxidation of catecholic compounds was expanded by the inclusion of 1,2-dihydroxy benzene (aka catechol; **(3)** in Figure 1) and 3,4-dihydroxy benzoic acid (**(4)** in Figure 1). In a first experiment, about 100mg GA, dissolved in 8mL water, was mixed with about 25mg **(1)**, **(2)**, **(3)** or **(4)**, dissolved in 2mL methanol prior to the addition to the GA solution to ensure the dissolution of the four compounds. All mixtures were kept at 37°C for multiple days. Fig. 7 shows a photograph taken of these mixtures after five days of reactions.

**Fig. 7:**
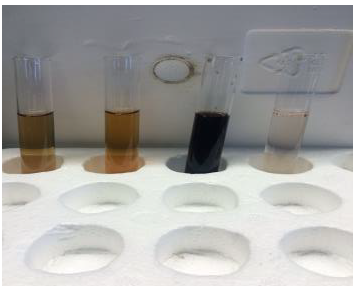
Photograph of reaction mixtures containing 100mg GA and 25mg, from left to right, **(1)**, **(2)**, **(3)** and **(4)** after five days of reaction at 37°C.

As the reaction involving **(4)** showed little formation of pigment, only the reaction mixtures involving **(1)**, **(2)** and were dialyzed and lyophilized.

In a second set of experiments, about 30mg of **(3)** was dissolved in 10mL water and placed inside the well of a cell culture dish. About 100mg of GA or other PS materials was added and the mixtures were kept at RT for six days. Fig.8 shows a photograph of this experiment after four days of reaction.

**Fig. 8:**
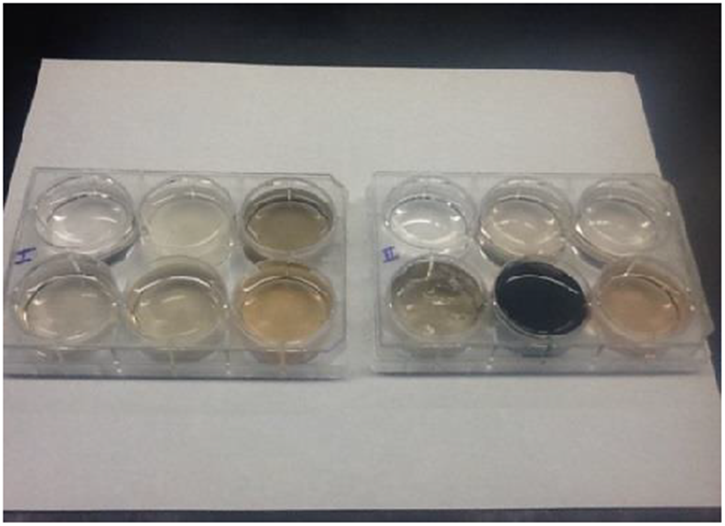
Photograph of reaction mixtures containing 30mg **(3)** and 100mg test substances after four days of reaction at RT. Test substances were: top row, from left to right, none – alignate – fucoidan – L-ascorbic acid – dextran sulfate – sodium glucuronate; bottom row, from left to right, carboxymethylcelulose – sodium acetate – chondroitin sulfate type C – carrageenan – gum Arabic – chondroitin sulfate type A.

The mixture involving GA and **(3)** was siphoned off, diluted for SEC analysis, dialyzed against water and lyophilized. Fig. 9 presents the SEC profile of an aliquot of this reaction between GA and **(3)**.

**Fig. 9:**
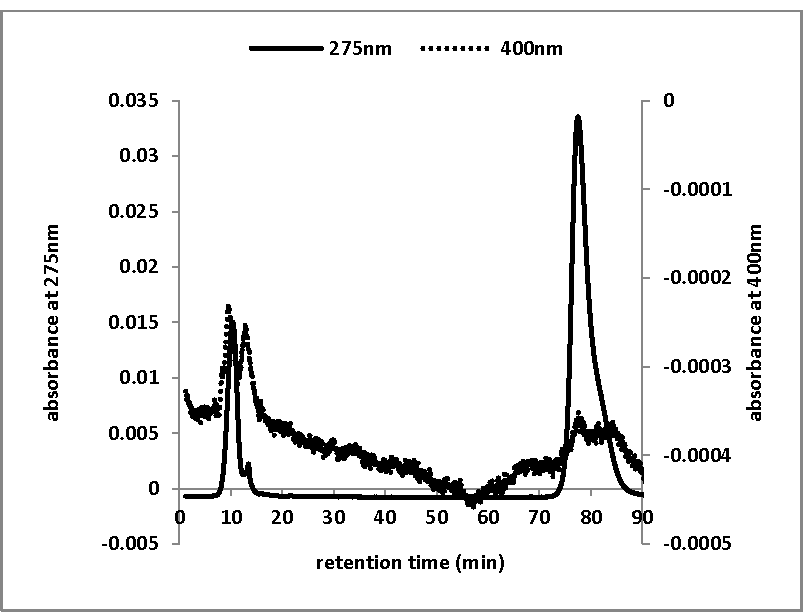
SEC profile of an aliquot from a reaction mixture containing GA and **(3)**. After four days of reaction, an aliquot of the reaction mixture containing 30mg **(3)** and 100mg GA was diluted 20-fold and analyzed by SEC.

When viewed at 250nm, peaks in region II of the SEC profile can be observed and a dominant peak with a retention time of about 80 minutes, associated with **(3)**, in region IV of the profile. When viewed at 400nm, peaks in region II of the SEC profile can be observed, but they are barely visible above the baseline. Despite the intense black color of the reaction mixture (see Fig. 8), it contains a significant amount of unreacted **(3**) and a minimal amount of darkly-colored pigment.

Fig. 10 represents FT-IR scans of GA and the purified products of the reaction, shown in Supplemental Figs. 1 or 2, between GA and **(1)**, **(2)** or **(3)**.

**Fig. 10:**
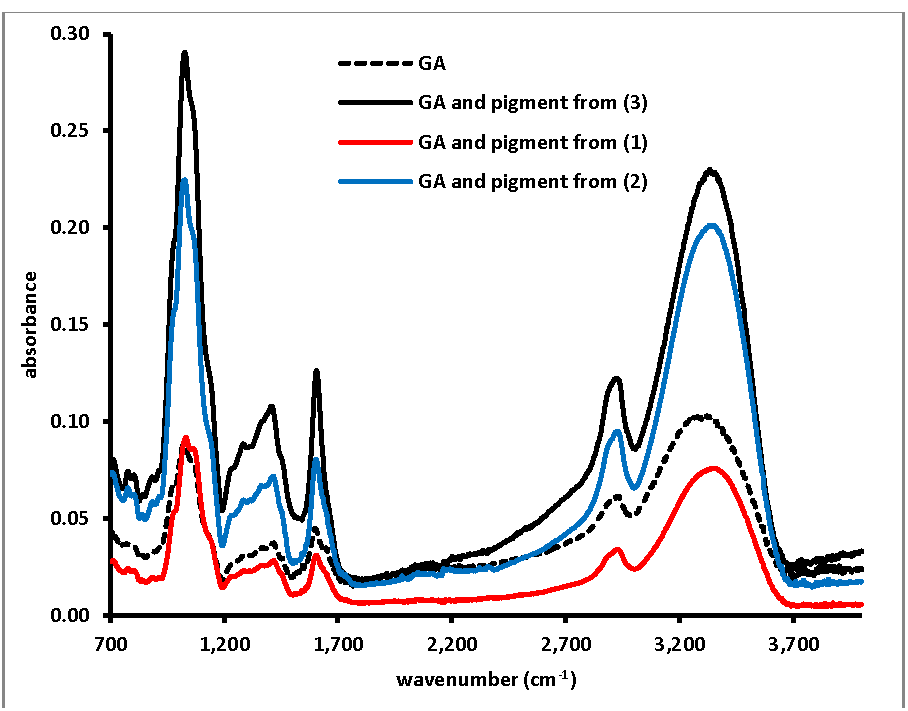
FT-IR spectra of pigment materials from the reactions between GA and **(1)**, **(2)** or **(3)**. FT-IR spectra of the purified and lyophilized pigments obtained from the reactions between GA and **(1)**, **(2)** or **(3)** as shown in Supplemental Figs. 1 or 2.

As for our observations regarding the reactions between select PS and CAs, the FT-IR scans of the GA-pigment products seem qualitatively similar to the FT-IR spectrum of GA. Despite the obvious physical differences between the materials, GA is a very light-brown colored powder, while the GA-pigment preparations are gold- (pigments made from **(1)** or **(2)**) or black-colored (pigment made from **(3)**), there appear to be very little chemical differences between all the materials as judged from FT-IR spectroscopy. On a weight-by-weight basis, the GA-pigment materials consist primarily of GA with very little contribution from the pigment fraction present.

### 3.3. Ester hydrolysis by GA

As described in the results from the RP-HPLC and SEC analyses of the interaction between GA and **(2)**, no evidence of the release of **(1)** from **(2)** could be observed. However, the lack of **(1)** could be due to the fact that it may be oxidized by GA immediately upon hydrolysis from **(2)**. Thus the capacity of GA to hydrolyze esters was evaluated by mixing GA with **(6)** and monitoring the decline in **(6)** and increase in **(5)** through RP-HPLC analyses.

Mixtures were set up containing **(6)** (1.0mM) with or without GA (tested between 2 and 18mg/mL) at 37°C and analyzed by RP-HPLC after 24 hours. The AUC (signal at 260nm) for both **(6)** and **(5)** was determined and plotted as a function of GA concentration in Fig. 11.

**Fig. 11:**
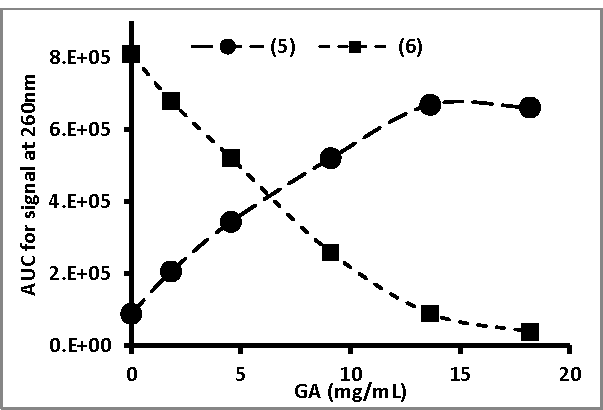
Effect of GA concentration on the hydrolysis of **(6)** to **(5)**. GA, at varying concentrations, was incubated with 1mM **(6)** at 37°C and after 24h aliquots of the reaction mixtures were analyzed by RP-HPLC and the AUC (signal at 250nm) for both **(6)** and **(5)** was determined.

The pattern of results indicated that GA enhanced the hydrolysis of **(6)** to **(5)** in a concentration-dependent fashion. In a second set of experiments, **(6)** (1 mM) was mixed with GA (9 mg/mL) that was kept at RT or kept at 60°C for multiple hours prior to incubation with **(6)**. All experiments were set up in triplicate and Fig.12 present the AUC (as viewed at 260nm; average ± standard deviation) of both **(6)** and **(5)** after 6h and 25h of incubation at 37°C with: a) water, b) GA solution kept at RT and c) GA solution kept at 60°C for multiple hours prior to incubation with **(6)**.

**Fig. 12:**
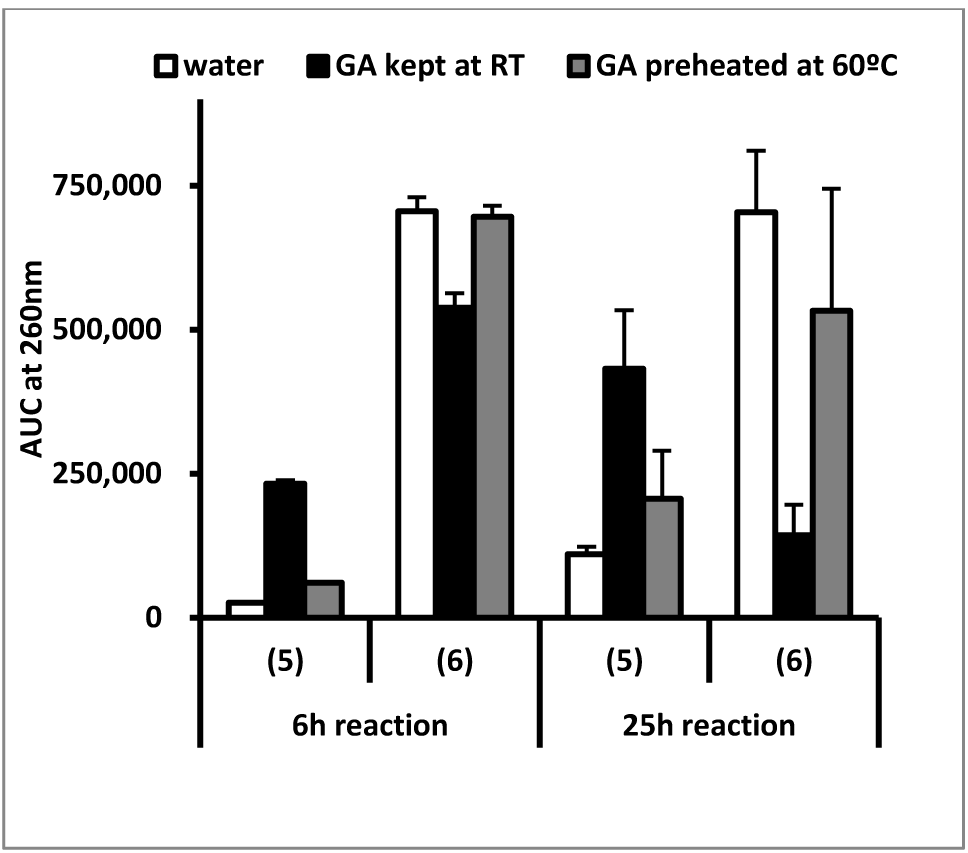
Effect of GA, kept at RT or preheated at 60°C, on the hydrolysis of **(6)** to **(5)**. GA, at 9 mg/mL and either kept at RT or preheated at 60°C for multiple hours, was incubated with 1mM **(6)** at 37°C. After 6 and 25h, aliquots of the reaction mixtures were analyzed by RP-HPLC and the AUC (signal at 260nm) for both **(6)** and **(5)** was determined. All experiments were performed in triplicate and the bars represent the average AUC ± standard deviation.

The results show that the GA solution kept at RT enhanced the hydrolysis of **(6)** when analyzed after 6h of reaction and significantly more when analyzed after 25h of reaction. The GA solution kept at 60°C had no or little significant effect on the hydrolysis of **(6)** compared to the incubation of **(6)** with water.

## 4. Discussion

The oxidation of **(1)** or **(2)** in the presence of transition metal ions has been studied before.[10, 11] Our study of the oxidation of **(1)** or **(2)** fits with our ongoing experiments regarding the PS-mediated oxidation of CAs and other compounds.[4] In that earlier report, the effect of GA on the oxidation of CAs was only briefly touched upon and was not included in that report. However, some preliminary results indicated that GA was capable of promoting the auto-oxidation of CAs, but not as intensely as for other PS like the different types of CS or FUCO.^2^ On the other hand, the auto- or Cu^2+^-mediated oxidation of **(1)** by the PS mentioned in our earlier report was investigated. In general, these other PS materials did not, or only weakly, promote the auto-oxidation of **(1)**. However, nearly all PS materials tested did promote the Cu^2+^-mediated oxidation of **(1)**. ^3^ Thus, preliminary observations had indicated that GA was capable of promoting the oxidation of **(1)** in the absence of any transition metals, while it appeared to have a much milder effect on the auto-oxidation of CAs compared to other PS materials.

The results presented in Fig.2 indicated that GA promotes the oxidation of **(1)**, independent of the presence of any redox-sensitive ions. In addition, the results presented in Fig.3 indicate that this effect of GA is concentration dependent. The chromatographic studies (RP-HPLC and SEC) confirmed the observations from the UV-Vis studies and indicated that, in the presence of GA, both **(1)** and **(2)** are converted into high molecular mass, pigmented materials (see Figs 5 and 6). In addition, GA appeared unique among PS-based materials in its capacity to promote the oxidation of **(3)** into a high molecular mass, pigmented materials (see Figs. 8 and 9). It is worth noting that GA appeared not to promote the oxidation of **(4)** into pigmented material (see Fig. 7). The action of GA on phenolic compounds appears to exhibit some substrate selectivity, echoing some of the results of previous research on the pro-oxidizing capacity of GA materials.[12] Although GA appears to have a similar reactivity towards **(1)** and **(2)**, the SEC profiles of the reaction mixture containing GA and **(2)** appeared to contain more peaks in its profile (see Fig. 6) compared to the SEC profile of the reaction mixture containing GA and **(1)** (see Fig. 5). In our study of the oxidation of **(1)** by GA we did not observe any new peaks in the chromatographic analyses other than the sharp peak in region I of the SEC analyses (see Fig. 5, panels A and B). The SEC retention time of this peak suggests it corresponds to a material with very high molecular mass. This could be an aggregate of multiple high molecular mass molecules including the oxidation products of **(1)** or this could be a nanoparticle type of material stabilized by the components of GA. SEC analyses of the oxidation of **(2)** did show the emergence of a new peak with significant absorbance in the VIS range of the spectrum in region III of the SEC profiles (see Fig. 6). Only later in the reaction was a sharp peak in region I of the SEC profiles observed; similar as observed for the reaction with **(1)**. Similarly, the SEC profile of the reaction mixture containing GA and **(3)** (see Fig. 9) contains a limited number of peaks. These comparisons may indicate that the reactivity between GA and **(2**) follows a more complex pattern; a pattern whereby more intermediates are generated during the reaction between GA and **(2)**.

Despite the visually dramatic effects of the reactivity between GA and **(1)**, **(2)** or **(3)** (see Figs. 7 and 8), the purified GA/pigment material appears to contain mostly GA and very little pigment as judged from the FT-IR spectroscopic results (see Fig. 10) Similar observations were made for the interaction between PS and CAs.[4] For the synthesis of MN-like pigments from DOPA in the presence of select PS, the contribution of the pigment fraction to the PS/pigment material has been estimated to be between 1.4 and 2.4 % (w/w).[13]

The capacity of GA to promote the auto-oxidation of compounds like **(1)**, **(2)** or **(3)** may be due to the fact that this material can be expected to contain redox sensitive elements like Cu and particularly Fe.[14] However, as mentioned earlier, GA had only a modest effect on the auto-oxidation of CAs and these compounds are readily oxidized in the presence of redox ions like Cu^2+^ or Fe^2+^. Deiana et al. studied the oxidation of **(1)** by Fe^3+^ in a calcium- polygalacturonate network.[10] They observed the formation of an absorbance band around 680nm in the UV_Vis spectrum, but this was attributed to the formation of a physic-chemical complex between **(1)** and Fe^3+^. In our experiments, when GA was mixed with **(1)**, no immediate change in color, nor an absorbance band around 680nm, could be observed. Thus, we believe the capacity of GA to auto-oxidize select catecholic compounds is due to other (than redox-sensitive cations) chemicals present in the GA material. Billaud et al. described the partial purification of a PPO from GA.[12] This enzyme appeared to be stable when stored in solution with pH 4.5 at 20°C for over 40 days. These results are in line with our observation of the thermal stability of the pro-oxidizing capacity of GA following its incubation at 60°C for multiple hours. However, no follow-up studies on this PPO from GA appeared to have been reported. PPOs are an important class of enzymes as they are almost universally present in animals, plants, fungi and bacteria.[15] Despite extensive research on this class of enzymes, there appears to be a significant lack of understanding between the presence of this type of enzyme and its precise role in, e.g., browning reactions.[15] The chemistry of GA is complex but consists mostly of PS type of materials. Up to 14% of the GA material may consist of glucuronic acid.[1, 14] As argued for the PS-mediated oxidation of CAs, the anionic PS fraction of GA, possibly chelating copper or iron ions naturally present in the GA material, could provide an environment that promotes the oxidation of **(1)**, **(2)**, **(3)** and other compounds. Thus, we would like to raise the hypothesis that the PS fraction present in GA is responsible for the pro-oxidizing properties observed for GA.

Despite the capacity of GA to hydrolyze esters like **(6)**, in both our RP-HPLC and SEC analyses we did not observe any release of **(1)** from **(2)** when the latter was mixed with GA. Either no hydrolysis of **(2)** occurred and GA directly oxidized **(2**) or any **(1)** released through hydrolysis of **(2)** by GA was immediately oxidized further. It is worth noting that despite the thermal stability of the oxidizing capacity of GA, the ester hydrolysis capacity of GA appeared to be thermally unstable (see Fig. 12) suggesting that these two chemical activities are associated with different components of the GA material.

In general, both MNs and lignins are pigmented polymers of which the monomeric building blocks are ultimately derived from phenylalanine through the action of various enzymes, including PPOs for plant-derived MNs.[5, 8, 16] Despite the extensive body of work that exists on the topic of MNs and lignins, there appears to be some uncertainty regarding the ultimate mechanism of the polymerization of the phenyalanine-derived monomers into the pigmented, macromolecular structures.[5, 8, 17] It has been suggested that the polymerization into the pigmented substances is a secondary event relative to the generation of the monomeric substances. In other words, independent events may determine the type and amount of monomers that are available for polymerization; the polymerization itself may depend on totally different factors.[15, 18] In this context it is important to note that the hydrolysis of **(2)** has recently been recognized as a very important step in the biosynthesis of lignins.[9] Our observations described in this work indicate that materials like GA may be capable of creating high molecular mass materials out of diphenolic monomeric components. This is in line with our observations regarding the generation of high molecular mass, pigmented, materials derived from CAs in the presence of PS. All these observations leads us to hypothesize that PS, ubiquitously present in the extracellular environment, may be important factors in promoting or guiding the formation of high molecular mass, pigmented, materials derived from diphenolic compounds.

## Acknowledgements

The research and Lauren Tyler were supported by a grant (#P031B090214) from the US Department of Education. Jade Readus was supported by a grant (#2T34GM007663) from the National Institutes of Health (NIH).

## Footnotes

1 Personal communication

2 Personal communication

3 Personal communication

